# A scaleable inducible knockout system for studying essential gene function in the malaria parasite

**DOI:** 10.1101/2024.01.14.575607

**Authors:** Abhinay Ramaprasad, Michael J Blackman

## Abstract

The malaria parasite needs nearly half of its genes to propagate normally within red blood cells. Inducible ways to interfere with gene expression like the DiCre-lox system is necessary to study the function of these essential genes. However, the existing DiCre-lox strategy is not well-suited to be deployed at scale to study several genes simultaneously. To overcome this, we have developed SHIFTiKO (frameshift-based trackable inducible knockout), a novel scaleable strategy that uses short, easy-to-construct, barcoded repair templates to insert *loxP* sites around short regions in the target genes. Induced DiCre-mediated excision of the flanked region causes a frameshift mutation resulting in genetic ablation of gene function. Dual DNA barcodes inserted into each mutant enables verification of successful modification and induced excision at each locus and collective phenotyping of the mutants, not only across multiple replication cycles to assess growth fitness but also within a single cycle to identify the specific phenotypic impairment they exhibit. As a proof of concept, we have applied SHIFTiKO to screen the functions of malarial rhomboid proteases, successfully identifying their blood stage-specific essentiality. SHIFTiKO, thus offers a powerful platform to conduct inducible phenotypic screens to study essential gene function at scale in the malaria parasite.

## Introduction

The ability to manipulate genes in the malaria-causing parasite, *Plasmodium*, plays a major role in efforts to study the biology of this parasite in the laboratory (*de Koning-Ward et al., 2015*). This remains highly relevant as malaria continues to be a major global health problem causing 608,000 deaths in 2022 (*Organization, 2023*). Emerging resistance to frontline antimalarial drugs further threatens our concerted efforts to reduce the malaria disease burden. Over 35% of the ∼5000 protein-coding genes in the genome of this early-diverging unusual eukaryote lack any known functional domains (*Oberstaller et al., 2021*) that might inform on their putative function. As a result, progress in our overall understanding of *Plasmodium* biology relies upon the pace at which researchers can interrogate and discover gene function in this parasite (*Ishizaki et al., 2022*).

Systematic study of gene function is challenging in *Plasmodium*. Despite work over the past decade successfully adapting CRISPR-Cas9 technology in the parasite (*Ghorbal et al., 2014*; *Wagner et al., 2014*), the lack of canonical non-homologous end-joining machinery (*Kirkman et al., 2014*) and the suboptimal efficiency of integrating exogenous DNA into the genome through homologous recombination still hinders our capability to perform gene disruption at scale. These challenges notwithstanding, immense efforts have screened gene essentiality at the genome scale using gene targeting vector libraries in the rodent malaria parasite *P. berghei* (*Bushell et al., 2017*) and non-targeted transposon mutagenesis in the most lethal human malaria parasite *P. falciparum* (*Zhang et al., 2018*). These screens have revealed that the parasite requires nearly half of its genes to propagate normally during the clinically relevant asexual blood stages, which involves successive cycles of in-traerythocytic replication. Conditional gene manipulation is necessary to study these essential genes (*Kudyba et al., 2021*). Malaria parasites are haploid during the blood stages and so disruption of an essential gene renders the mutant non-viable for further study. To address this, several inducible strategies have been implemented in *P. falciparum* to precisely regulate gene expression by inducing changes at the DNA (*Collins et al., 2013*), RNA (*Prommana et al., 2013*; *Ganesan et al., 2016*) or protein (*Armstrong and Goldberg, 2007*; *Muralidharan et al., 2011*; *Birnbaum et al., 2017*) level through the addition or removal of regulatory small molecules. Since parasite viability is not affected until the point of induction, these strategies allow an essential gene to be inactivated in a population of parasites at a desired time point and the resulting phenotype to be subsequently monitored.

One such inducible approach is the Dimerisable Cre (DiCre)-lox system in which a gene or critical sequence thereof can be inducibly deleted from the genome to disrupt its function using the site-specific recombinase Cre. Inducible control of Cre activity is brought about by constitutively expressing Cre in the parasite as two separate, enzymatically inactive polypeptides, respectively fused to rapamycin-binding FKBP12 and FKB proteins. A target region containing the gene of interest is marked for removal by flanking it with short, 34 bp *loxP* sequences that are recognisable by Cre. Addition of the drug rapamycin (RAP) dimerises the two Cre subunits to restore recombinase activity, and the active DiCre excises the *loxP*-flanked (‘floxed’) sequence. This process is highly efficient, capable of excising the target locus in close to 100% of parasites in a population upon treatment with RAP and can result in complete ablation of the gene’s expression to produce a definitive observable phenotype within the same or subsequent erythrocytic cycle (*Collins et al., 2013*; *Knuepfer et al., 2017*). In important refinements of this technology, artificial *loxP*-containing intron modules (*loxPint*) (*Jones et al., 2016*) that can be used to silently introduce *loxP* sites within an open reading frame and the use of CRISPR-Cas9 to flox sequences (*Knuepfer et al., 2017*) now allows the selective targeting of any region within any gene to rapidly generate inducible knockout mutants in *P. falciparum*. These combined strategies have been highly effective in studying genes with indispensable roles in parasite development (*Burda et al., 2023*; *Ramaprasad et al., 2022*), egress (*Thomas et al., 2018*; *Nofal et al., 2021*), invasion (*Patel et al., 2019*; *Perrin et al., 2021*) and pathogenesis (*Davies et al., 2020*; *Subudhi et al., 2023*) during blood stages and sensitive enough to discern even mild phenotypic defects in the case of some non-essential genes (*Pietsch et al., 2023*; *Ramaprasad et al., 2023*).

Whilst useful for detailed functional analysis of individual genes, the current workflow is not scaleable for functional screening of larger cohorts of genes for a number of reasons. To flox a region within a gene, it must be replaced with a ‘recodonised’ version incorporating silent base substitutions in order to retain gene function prior to excision and to protect from repeated Cas9-mediated cleavage at the target sites (*Knuepfer et al., 2017*). Commercial synthesis of these gene repair constructs at scale can be expensive and requires long turnaround times depending on the length of the floxed region. Suboptimal transfection efficiency in *P. falciparum* means that successfully modified parasite clones have to be isolated and expanded to a workable culture volume before gene excision can be induced, which can take several weeks (*Thomas et al., 2016*). Once excision of the floxed region is confirmed by PCR, phenotyping must ensue through growth assays that measure mutant growth fitness and phenotypic assays that reveal the nature of the limiting phenotypic defect using microscopy and flow cytometry. These experiments require extensive time, effort and reagents, which can be prohibitive for targeting multiple genes. Whilst the use of selection-linked integration (SLI) (*Birnbaum et al., 2017*), an approach that improves the rates of obtaining modified lines in *P. falciparum*, has enabled systematic generation of several inducible knockout lines (*Davies et al., 2020*; *Kimmel et al., 2023*), costly synthesis of recodonised regions and resource-intensive phenotyping still limits the use of this approach. Therefore, the current DiCrelox strategy must be reimagined if we are to perform inducible gene knockouts at scale in the malaria parasite.

Here, we have developed a frame<u>shift</u>-based <u>t</u>rackable <u>i</u>nducible <u>k</u>nock<u>o</u>ut (SHIFTiKO) system that uses short, modular repair templates to flox small regions within genes and simultaneously tag them with DNA barcodes. Induced excision introduces a frameshift mutation that renders the gene non-functional and the incorporated barcodes enable phenotyping of several mutants in a pooled manner. We demonstrate the utility of SHIFTiKO as a scaleable refinement to the DiCre-lox system by screening the functions of a family of rhomboid proteases in *P. falciparum*.

## Results

### SHIFTiKO design

First, we sought a floxing strategy that can be uniformly applied across genes irrespective of their size, thereby removing the need to synthesize long repair templates. We chose a recently adopted (*Ramaprasad et al., 2023*; *Bahl et al., 2021*; *Sherling et al., 2019*) frameshift-based gene disruption approach where a short segment within the target gene open reading frame, upstream of any putative functional domains, and of a length not divisible by three is floxed by inserting two closely-opposed loxP-containing intron (*loxPint*) modules. DiCre-mediated excision of the floxed region is predicted to result in a translational frameshift and the introduction of multiple downstream premature stop codons, thus truncating the gene and rendering it non-functional (Figure 1). Next, we aimed to tag the modified gene with a DNA barcode so that abundance of each parasite mutant can be quantified at the molecular level, thereby enabling simultaneous monitoring of the fitness of several mutants within uncloned populations. For this, a unique 12 base pair (bp) barcode was inserted next to the *loxP* site in two heterologous *loxPint* sequences (SERA2:*loxPint* (*Jones et al., 2016*) and SUB2:*loxPint* (*Collins et al., 2020*)) to create a pair of barcoded loxP-containing introns (or *boxit*; Figure 1). Specific primer-binding sites for barcode amplification were also added in such a way that a 77 bp long amplicon will contain the barcode from the first flanking *boxit* (*boxit*-) pre-excision and the barcode from the second flanking *boxit* (*boxit+*) post-excision of the floxed region. Finally, a new dual-guide targeting plasmid (pCas9-Duo) was generated that expresses two distinct guide RNAs (gRNA) in order to enhance the chances of a successful Cas9-mediated modification at the target region (Supplementary Figure S1A). Presently, this failsafe is achieved by multiple transfections, each using different single-guide targeting plasmids.

**Figure. 1.**
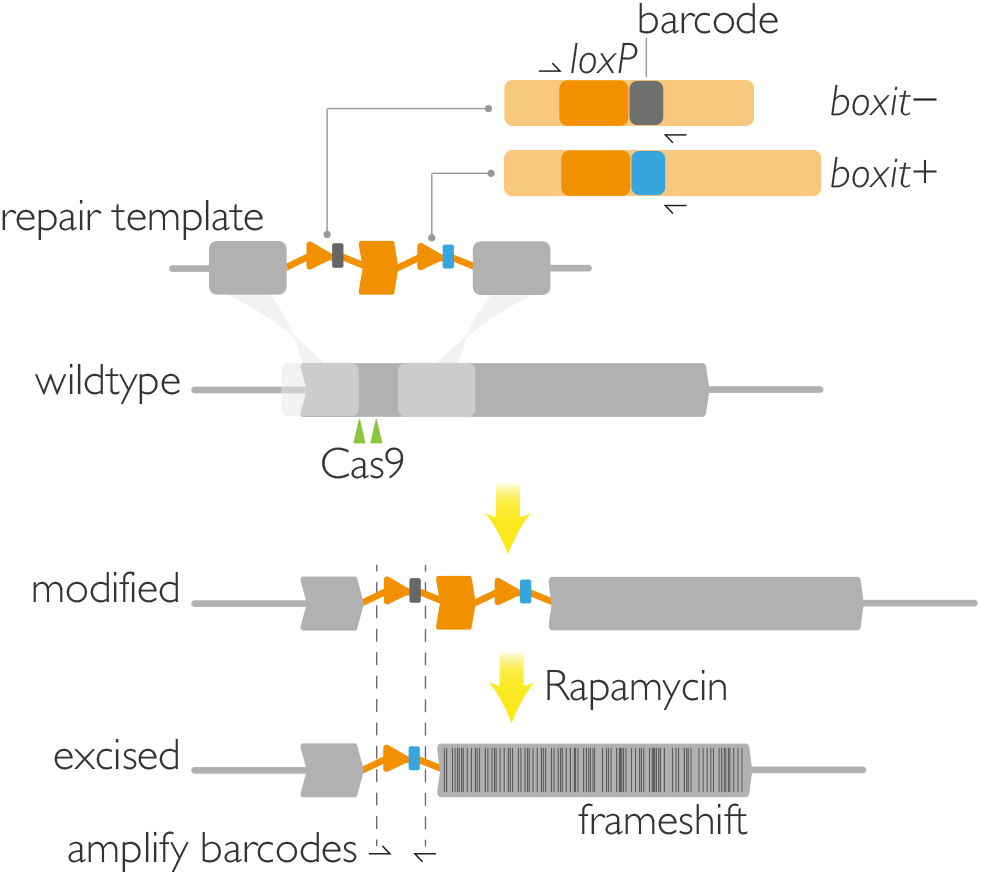
SHIFTiKO design. A short segment of length not divisible by three is replaced in the target gene with a recodonised sequence (orange block) flanked by a pair of synthetic intron modules (*boxit* ; orange line), each containing a *loxP* site (orange arrow head) and a unique 12 bp barcode. Gene modification is achieved by a Cas9-driven homology directed repair strategy in which double-stranded breaks are introduced in two gRNA target sites (green) simultaneously. Treatment with rapamycin induces DiCre-mediated excision of the segment, resulting in a frameshift mutation to render the downstream segment of the gene non-functional. Binding sites for barcode-amplifying primers (black half arrows) are placed in such a way that only the *boxit-* barcode (grey) is amplified from a modified *shiftiko* line and only the *boxit+* barcode (blue) is amplified from the excised mutant.

### SHIFTiKO creates barcode-trackable inducible mutants

To test our strategy, we targeted the *sub1* gene (PF3D7_0507500), which encodes a serine protease essential for egress of blood stage parasites (*Thomas et al., 2018*). To do this, a 199 bp segment upstream of the subtilisin-like catalytic domain was floxed with *boxit* modules. Treatment with RAP resulted in efficient DiCre-mediated excision of the floxed sequence (Figure 2A) and the loss of SUB1 expression, which as expected proved lethal to the parasite (Figure 2B) due to a severe egress defect (Figure 2C). Interestingly, diagnostic PCR of the entire target locus, whilst amplifying the modified locus (2,647 bp in - RAP, Figure 2A), did not produce a smaller amplicon (2,303 bp) from the unmodified locus which is generally expected when sampling transfectant populations that consist of both modified and wildtype parasites. This suggests a high rate of gene modification, possibly a result of expressing two gRNAs simultaneously and using shorter repair templates in SHIFTiKO. The transfectant parasites phenocopied clonal SUB1 iKO lines (*Thomas et al., 2018*) upon treatment with RAP indicating that SHIFTiKO can produce near-clonal populations of inducible mutants suitable for direct phenotyping.

**Figure. 2.**
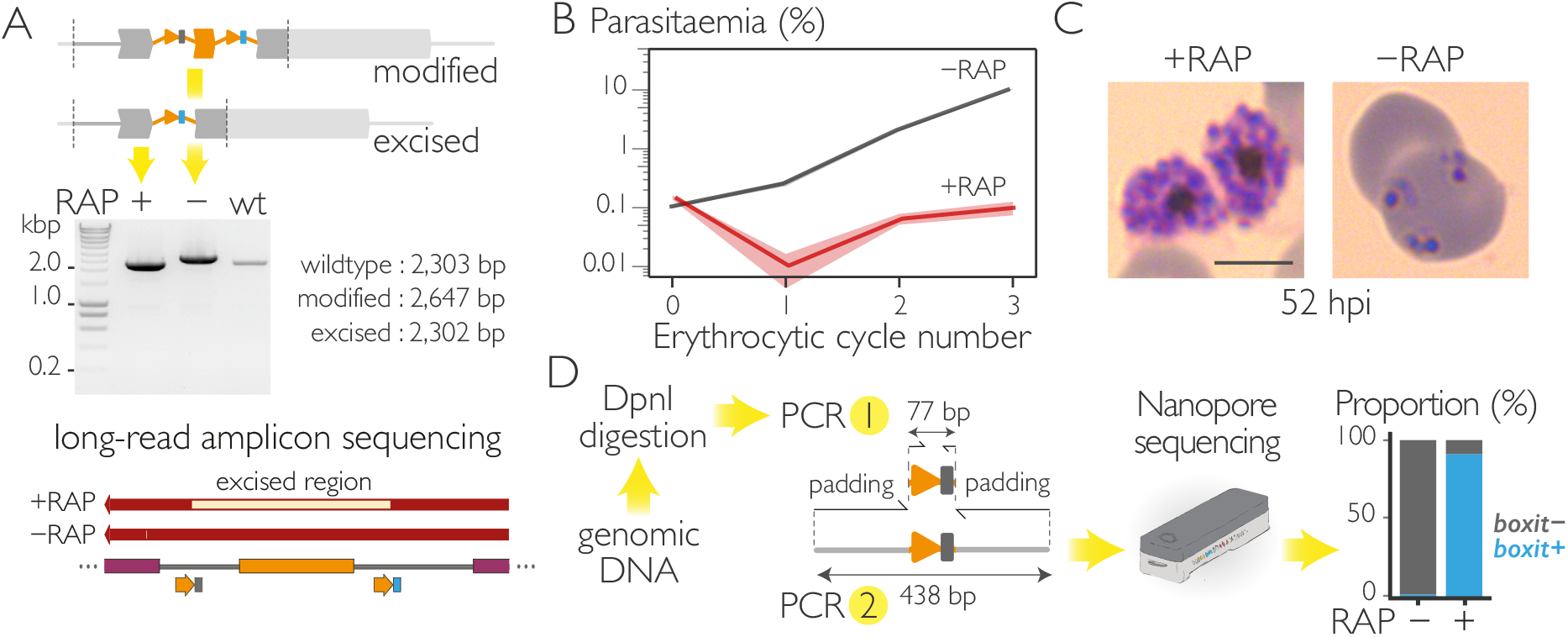
Induced disruption of SUB1 function using SHIFTiKO. (A) Diagnostic PCR amplification of the entire modified locus from *sub1-shiftiko* parasites at 24 h following RAP or mock treatment confirms efficient excision. Expected amplicon sizes are indicated. Long-read sequencing of the amplicons show precise excision of the floxed region containing the *boxit-* barcode whilst the *boxit+* barcode is retained in the RAP-treated parasites. (B) Treatment with RAP results in ablation of growth in an uncloned transfectant population of *sub1-shiftiko* parasites. Data shown are averages from three biological replicates using different blood sources (shaded ribbon, ± SEM). (C) Light microscopic images of Giemsa-stained *sub1-shiftiko* parasites at 52 h following RAP and mock treatment at ring stages show that RAP-treated mutants are unable to undergo egress. Scale bar, 5 μm. (D) To amplify barcodes from RAP- and mock-treated *sub1-shiftiko* parasites, genomic DNA extracts were first digested with DpnI to selectively remove residual repair plasmids carried over from the transfectant cultures. Nested PCR results in 438 bp long amplicons that were sequenced on a ONT MinION device. Barcode counts show increase in *boxit+* proportions in the RAP-treated population signifying efficient excision.

Next, we wanted to test whether the incorporated barcodes can be used to quantify the mutants, which in this case means distinguishing between excised and unexcised parasites using the *boxit+* and *boxit-* barcodes respectively. Barcodes were amplified from genomic DNA extracted from RAP- and mock-treated *sub1-shiftiko* parasites at 24 hours post-treatment using a generic pair of primers (barcode.F and barcode.R), yielding a 77 bp product. To explore the use of nanopore sequencing (Oxford Nanopore Technologies or ONT) to analyse these barcodes, we further performed nested PCR to add a random padding sequence on either side of the amplicon, thereby increasing its size to 438 bp to meet ONT’s minimum fragment size specifications. This step can now possibly be skipped with the new “Short Fragment Mode” configuration (released (*Technologies*, *2022*) during the course of this study) that allows sequencing of fragments as short as 20 bp. Barcode-sequencing (bar-seq) clearly showed a rise in *boxit+* barcodes in the RAP-treated population, measuring an excision rate of 91% at the time point of sampling (Figure 2D). Taken together, these results show that *boxit* modules can be efficiently inserted into a gene without affecting its function to prime it for an effective knockout by induced frameshift mutation and the proportion of mutants in the resulting parasite population can be quantified at any point by bar-seq.

### SHIFTiKO enables inducible phenotypic screens in *P. falciparum*

Encouraged by the above results, we wanted to explore whether several *shiftiko* mutants can be phenotyped simultaneously by tracking their barcodes in a mixed population. As the test gene cohort for this work, we chose to target a family of *Plasmodium* genes encoding rhomboids, intramembrane proteases that cleave membrane-bound substrates and that are known to play important roles in protozoan parasites (*Deu, 2017*; *M*. *Santos et al., 2012*). Whilst previous work has suggested that four to five of the eight *Plasmodium* rhomboids are likely essential (*Zhang et al., 2018*; *Bushell et al., 2017*; *Lin et al., 2013*) (Supplementary Figure S2), a systematic study to identify the precise biological functions of these putative enzymes is lacking. To ensure that the approach used was capable of discriminating a range of growth phenotypes, we included in the analysis additional well-characterised genes displaying four essential knockout phenotypes previously observed with the DiCre system, namely a limiting defect during trophozoite development (*gdpd* (*Ramaprasad et al., 2022*); PF3D7_1406300), schizont development (*pi-plc* (*Burda et al., 2023*); PF3D7_1013500), egress (*sub1* (*Thomas et al., 2018*); PF3D7_0507500) or invasion (*gap45* (*Perrin et al., 2018*); PF3D7_1222700), and the non-essential *p230p* (*van Dijk et al., 2010*) (PF3D7_0208900) as controls in our screen. Uniquely barcoded *boxit* modules were inserted into the 13 target genes to flox a region of length varying from 80 to 227 bp (Figure 3A). In the case of the rhomboid genes, we also explored the possibility of simultaneously tagging the N-terminus of each protein with a triple-hemagluttinin (3xHA) epitope to aid downstream detection of the protein. To do this, we modified our strategy slightly to include a 3xHA sequence and a second recodonised sequence to replace the region extending from the start codon to the segment to be floxed (Figure 3A). After transfecting parasites with SHIFTiKO plasmids and selecting them using the antifolate drug WR99210 (WR), we observed WR-resistant parasites in 9 out of the 13 cultures within 2-3 weeks. For the transfections that did not generate drug-resistant parasites, we hypothesized that tagging the N-terminus might be deleterious in the case of these genes (*rom4, rom6, rom7* and *rom10*) and subsequently were able to acquire drug-resistant parasites using repair constructs that did not incorporate the 3xHA tag.

**Figure. 3.**
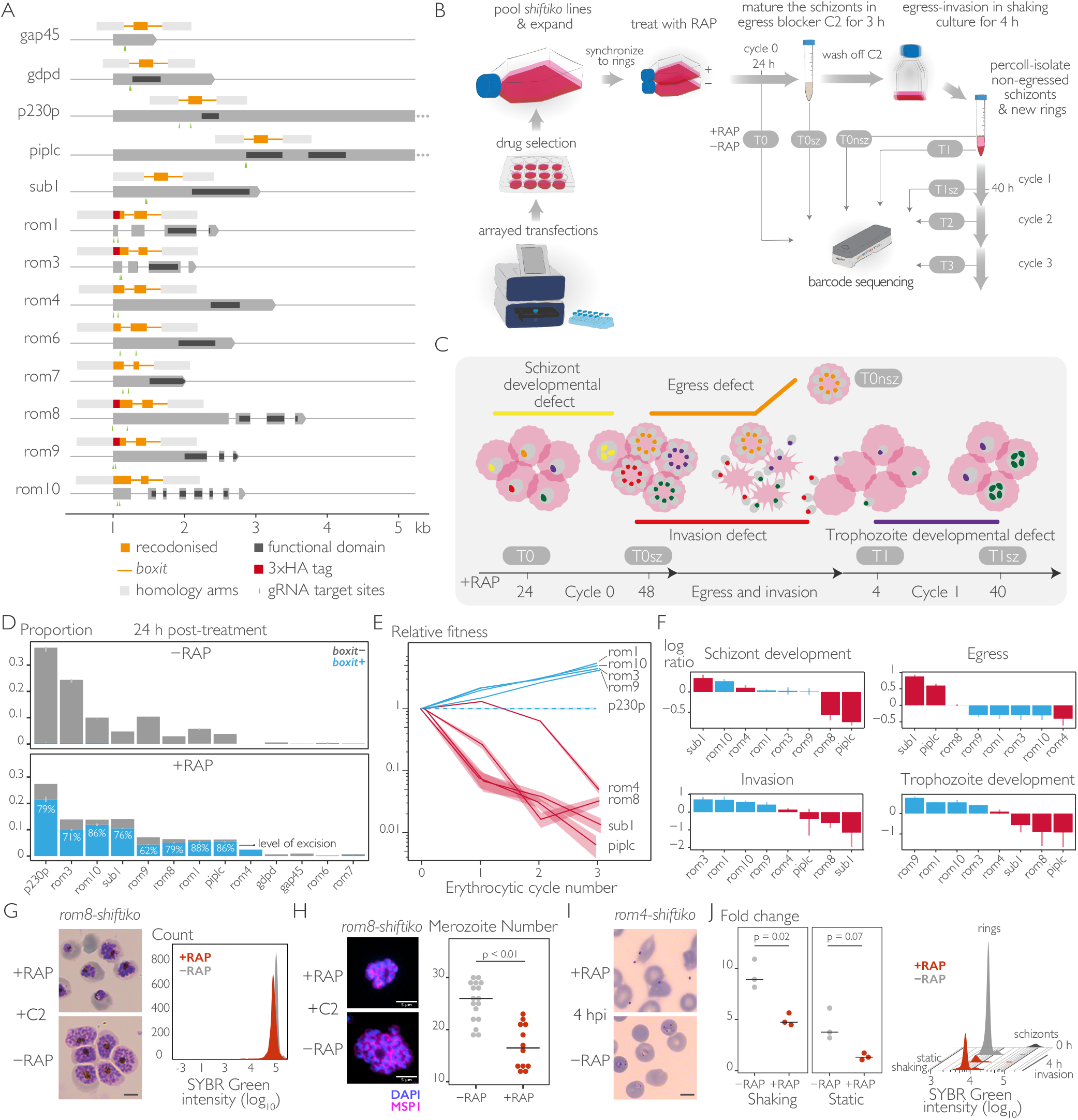
Inducible phenotypic screening of *Plasmodium* rhomboids using SHIFTiKO. (A) Uniform floxing strategies to create *shiftiko* lines for the 13 target genes (for gene IDs, see Supplementary Figure S2). In each case, a short region upstream of the functional domain and containing two good scoring gRNA target sites was chosen to be floxed with *boxit* modules. Simultaneous tagging of the N-terminus with a 3xHA tag was successful for four of the eight rhomboids attempted. (B) SHIFTiKO workflow to assess fate of several inducible mutants during blood stage progression. Transfections and subsequent drug-selection for transfected parasites are carried out in arrayed format to produce individual *shiftiko* lines, that are then pooled together, expanded and synchronized to set up a culture of tightly synchronized ring stages. Following rapamycin (RAP) treatment, samples are collected at carefully chosen time points during four successive erythrocytic cycles to capture the informative changes in barcode abundances - Mock-(-RAP) and RAP-treated (+RAP) samples from 24 h post-treatment (T0) to inform on successful integration and excision; fully mature +RAP schizonts from cycle 0 (T0sz), newly formed rings (T1) and the remaining non-egressed schizonts (T0nsz) after letting +RAP schizonts from cycle 0 undergo egress and invade fresh RBCs, and +RAP schizonts from cycle 1 (T1sz) to inform on knockout phenotypes; and samples from cycle 2 and 3 (T2 and T3) in addition to T0 and T1 to inform on growth fitness. Barcodes are amplified from genomic DNA from the collected samples and sequenced on a ONT MinION device. (C) Rationale behind choosing sampling time points within cycle 0 and 1 to evaluate knockout phenotypes. Mutants with a defect in schizont (yellow) or trophozoite (purple) development are expected to have less DNA copies and therefore lower barcode abundances in T0sz and T1sz respectively, compared to non-defective mutants (green). Barcodes of mutants with an invasion defect (red) are expected to be depleted in newly formed ring fractions (T1) whilst those of mutants with an egress defect (orange) are selectively enriched in the non-egressed schizont fraction (T0nsz). (D) Proportions of *boxit-* and *boxit+* barcodes in mock-(-RAP) and RAP-treated (+RAP) parasites (T0) show successful integration and excision in 9 out of 13 target genes. Data shown are averages from three replicate RAP treatments (error bars, ± SEM). (E) Relative growth fitness of knockout mutants (changes in *boxit+* barcode proportions from T0, normalised to the non-essential *p230p* gene) reveal essential (red) and non-essential (blue) genes (shaded ribbon, 95% confidence interval of the ratio estimated using delta method). (F) Within-cycle changes in barcode proportions (log_2_ ratios normalised to the non-essential *p230p* gene) reveal knockout phenotypes for the mutants (red, essential gene, blue, non-essential). Inferring the phenotypes from the first instance of aberrant change in barcode abundances during cycle progression, data reveals a schizont developmental defect (T0sz vs. T0) in *rom8* and *piplc*, egress defect (T0nsz vs. T0sz) in *sub1* and invasion defect (T1 vs. T0sz) in *rom4* mutants. Data shown are averages from three replicate experiments (error bars, ± SEM). (G) Light microscopic images of Giemsa-stained *rom8-shiftiko* schizonts allowed to fully mature in the presence of reversible egress inhibitor C2 following RAP (+RAP) and mock (-RAP) treatment at ring stages. ROM8-null parasites exhibit defective development producing abnormal schizonts, as confirmed and quantified using flow cytometry to measure parasite DNA content. Fluorescence intensity of the SYBR Green-stained RAP-treated population (red) was detectably lower than that of the mock-treated population (grey). Scale bar, 5 μm. (H) MSP1 (merozoite surface protein 1) localization in ROM8-null schizonts (+RAP) show evidence of segmentation to produce individual merozoites, albeit in significantly lower numbers compared to mock-treated controls (-RAP) (crossbar represents median; n=12; Welch t-test with Bonferroni adjusted p-value). Scale bar, 5 μm. (I) Light microscopic images of Giemsa-stained RAP- and mock-treated *rom4-shiftiko* parasites after 4 h of invasion in static cultures. ROM4-null merozoites are unable to invade RBCs and form ring stages. Scale bar, 5 μm. (J) Fold change in parasitaemia after 4 h invasion of mock-(-RAP) and RAP-treated (+RAP) *rom4-shiftiko* parasites under shaking and static conditions. RAP-treated cultures show no increase in parasitaemia under static conditions but show a 5-fold increase in shaking cultures, albeit still lower than mock-treated controls (crossbar represents median, three replicate RAP treatments with different blood sources; individual points represent each replicate). Fluorescence intensity of SYBR Green-stained parasite populations pre-(0 h) and post-invasion (4 h) confirm that ROM4-null mutants undergo normal egress (causing the disappearance of the schizont peak) and that the increase in parasitaemia in shaking cultures is due to formation of ring stages.

Ring stage parasites from each culture were combined in equal proportions to create a unified pool that could be continuously cultured, stage synchronized and RAP-induced to produce a mixed mutant population (Figure 3B). By analysing the barcodes from this pool at specific timepoints, we could collectively track the presence and fate of each barcodetagged mutant following induced mutagenesis. Initially, barcodes identified in the mock- and RAP-treated pool 24 h post-treatment (T0) verified successful integration and later RAP-induced excision at all targets, except for *gdpd, gap45, rom6* and *rom7* (Figure 3D). These results were validated by diagnostic PCR performed on the individual uncloned *shiftiko* lines (Supplementary Figure S3) with unsuccessful (*gap45* and *rom6*) or low level (*rom7*) of integration or inexplicable deletions within *boxit+* in the case of *gdpd* as causes for failure. In the case of *rom4*, whilst its *boxit-* barcode could not be detected, we observed appearance of its *boxit+* barcode upon treatment with RAP. Overall, a 60-90% level of excision was observed at the target genes in the pool after 24 h of treatment with RAP that then reached 97-100% by the end of cycle 0 (Supplementary Figure S4). Subsequently, to profile the replication fitness of the successfully generated mutants, we tracked the changes in barcode abundances across different replication cycles (T0, T1, T2, and T3, corresponding to cycles 0, 1, 2, and 3 post-treatment, respectively). It was anticipated that barcodes linked to essential genes would diminish or disappear entirely due to a pro-liferation defect in the respective knockout mutants. This expectation was confirmed as barcodes associated with essential control genes *sub1* and *piplc* were significantly reduced whilst barcode levels were sustained for the non-essential gene control *p230p* (Figure 3E). Our analysis further identified *rom4* and *rom8* as essential rhomboids, exhibiting a similar diminishing trend in their barcodes. The outcomes of the growth fitness screen corresponded well with the growth profiles recorded for individual uncloned *shiftiko* lines upon treatment with RAP (Supplementary Figure S5), thus validating the approach. More-over, in the case of *piplc*, whilst the presence of unmodified parasites in the RAP-treated uncloned population (evident from wildtype locus amplified by diagnostic PCR, Supplementary Figure S3) caused an uncharacteristic increase in parasitaemia in the cell-based growth assay, the barcode-based growth pro-file using SHIFTiKO was able to precisely discern the severe growth defect in *piplc* mutants from the confounding wildtype background.

Next, within-cycle changes in barcode abundance caused naturally by DNA replication during schizogony were exploited to evaluate the effects of gene disruption on parasite development (Figure 3B, C). We predicted that a drop in barcode abundance between ∼24 h trophozoites (T0) and mature ∼48 h schizonts (T0sz) during cycle 0 would indicate a schizont developmental defect, whilst a similar decline between ∼4 h rings (T1) and ∼40 h schizonts (T1sz) during cycle 1 would indicate a trophozoite developmental defect. This prediction was supported by the case of *piplc*, a gene previously shown to be essential for schizont development, showing a ∼1.7 fold reduction in its barcode levels at schizont stages (Figure 3F, schizont development). The essential rhomboid, *rom8* also showed a similar reduction, indicating a defect in schizont development. This was further validated by detailed phenotyping of ROM8-null mutants which showed a defect in late schizont development that results in abnormal schizont forms with a lower number of merozoites compared to mock-treated controls (Figure 3G, H). Finally, we sampled barcodes after a 4 h invasion window at the end of cycle 0 to assess invasion and egress processes. Barcodes that were enriched in nonegressed schizonts isolated following the expected period of invasion (T0nsz *vs*. T0sz) would point to an egress defect, whilst those barcodes *exclusively* depleted in the newly formed rings (T1 *vs*. T0sz) would indicate an invasion defect. The trend observed with barcodes from *sub1* (enriched almost 2-fold in T0nsz and severely depleted in T1), a gene absolutely essential for egress, confirmed our hypothesis (Figure 3F, egress and invasion). The trends also showed that gains made by *rom4* barcodes in the ring fractions were low given the rate of their egress, suggesting an invasion phenotype for the *rom4* mutant. As confirmation, detailed phenotyping of ROM4-null parasites revealed a profound defect in their ability to invade RBCs (Figure 3I), that could be overcome to a certain extent by mechanical shaking during invasion (Figure 3J). Incidentally, this also explains the distinct growth fitness profile for *rom4*, with the slight gain in fitness in cycle 1 (T1) evidently a result of invasion being performed under shaking conditions and the downward trend later on due to subsequent invasions occurring in static cultures. In conclusion, these data show that both growth fitness and the knockout phenotype can be evaluated with great precision in mutants of multiple genes in a single experiment using SHIFTiKO, thus paving for the first time the way for inducible phenotypic screens in *P. falciparum*.

## Discussion

Conditional mutagenesis remains the only genetic strategy for the functional analysis of essential asexual blood stage genes in malaria parasites. DiCre-mediated gene disruption, which entirely eliminates protein expression, provides a more reliable phenotype than inducible knockdown methods that only reduce protein levels, especially for proteins with enzymatic activity (*Kudyba et al., 2021*). With SHIFTiKO, we have now greatly enhanced the scaleability of this powerful tool by dispensing of the need for synthesizing long artificial genes, limiting dilution cloning steps and repetitive phenotypic assays. We have shown here through extensive validation that SHIFTiKO can reliably identify successful floxing, successful excision, and subsequent growth fitness and specific knockout phenotype for several genes within a single experiment that is reproducible across independent RAP treatments and pooled cultures (results from another independently run experiment can be found in Supplementary Figure S6). This completely replaces several time-consuming steps in the traditional DiCre workflow such as diagnostic PCR for confirming integration and excision, and laborious cell-based growth and phenotypic assays.

In its current form, SHIFTiKO can be used to deploy DiCre at scale to conduct targeted inducible screens of moderately sized groups of essential genes, such as a protein family or enzymes in a metabolic pathway. Importantly, it is well-suited for targeting very large genes and proteins without identified functional domains, as the induced frameshift strategy operates independently of such information. Pooled phenotyping significantly reduces the time needed to determine at which stage in the parasite’s asexual replication cycle a target gene is essential. This allows rapid identification of genes of interest, allowing resources to be focused on efforts to characterize them in detail whilst minimizing time spent on gene targets that turn out to be dispensable (*Burda et al., 2023*). Such detailed functional analysis can be performed by reverting back to the original *shiftiko* line but would ideally require the protein to be tagged to provide insights into expression profiles, subcellular localization and biochemical analysis (e.g. by pull-down). Whilst our work has shown that concurrent N-terminal tagging is feasible when generating *shiftiko* lines, it is more practical to design simpler repair templates initially then introduce the tag later in a subsequent gene-editing step. This is facilitated by the presence of the negative selectable marker *yFCU* in the pCas9-Duo plasmid, which is episomally carried by all *shiftiko* lines.

The ability to maintain several *shiftiko* lines simultaneously as a single pooled culture owing to the conditional nature of gene disruption offers considerable robustness and convenience. During the course of this study, we have routinely cultured these pools for many months, subjected them to Percoll-based synchronization, cryopreserved and revived them, and added additional *shiftiko* lines to make newer versions of the pool without compromising usability. The barcode sequencing protocol has been specifically tailored for use with the nanopore platform as a swift, flexible and economical approach that can be readily implemented in any laboratory setting without relying on external sequencing facilities. In our experiments, barcode amplification and library preparation could be completed within a few hours and sequenced within just 6-8 hours using a MinION Flow Cell or the more cost-effective Flongle. Both the duration of the run and the choice of flow cell are adjustable based on the required number of reads, which varies depending on the complexity of the mutant pool.

The method however still suffers from certain limitations. We failed to modify four of our target genes, including genes (*gap45* and *gdpd*) that have been successfully disrupted out in the past using traditional DiCre methodology. This suggests some genes (or regions within them) may be refractory to introduction of *boxit* modules in tandem. Efforts are underway to flox a different target region in these genes. Designing the floxing strategy, synthesizing the ∼550 bp repair template and assembling both targeting and repair plasmids for each gene can still be time-intensive, thereby limiting the number of *shiftiko* lines we could produce at a time. We have partially addressed this by recently optimising one-pot assemblies to produce the plasmids in a rapid and modular fashion (Supplementary Figure S1B,C). Experimenting with shorter *boxit* modules and floxed regions, as well as moving to a single-plasmid geneediting strategy and automating floxing design using in silico tools such as GeneTargeter (*Cardenas et al., 2022*) can further simplify this process in the future and improve SHIFTiKO’s scaleability.

The creation of trackable inducible mutant lines for the first time in *P. falciparum* opens up exciting possibilities to scale up gene function discovery in this parasite. The mutant parasite pool could be exposed to various selective pressures such as drug or substrate challenges to rapidly identify genes, disruption of which confers a selective advantage under such conditions. Additionally, technologies such as single-cell RNA sequencing could be employed to study the transcriptional states of these mutants in a powerful way. SHIFTiKO could also be potentially useful to conduct inducible screens in other *P. falciparum* life stages (*Tibúrcio et al., 2019*), rodent malaria models (*Kent et al., 2018*) and protozoan parasites (*Hunt et al., 2019*) with an established DiCre system.

## Methods

### Repair template design and plasmid construction

To create *boxit-* and *boxit+* modules, sequence changes were introduced around the *loxP* site in two heterologous *loxPint* sequences, SERA2:*loxPint* (*Jones et al., 2016*) and SUB2:*loxPint* (*Collins et al., 2020*) reasoning that the region which already holds a 34 bp *loxP* sequence would be able to sustain a few more changes without disrupting intron function. Four basepairs were modified upstream of *loxP* site to produce the forward primer-binding site for barcode amplification (barcode.F) in *boxit-* alone, whilst a 12 bp unique barcode followed by a 19 bp reverse priming-binding site (barcode.R) were inserted after the *loxP* site in both *boxit-* and *boxit+*.

All repair templates in this study were designed as follows. Two high-quality gRNA sequences in close proximity (<200bp) to each other and upstream of a gene’s functional domain were chosen from gRNAs predicted using EuPaGDT (*Peng and Tarleton, 2015*). A region that spans the two Cas9 cleavage sites and of length not divisible by 3 was chosen as the target segment to be recodonised and floxed (FR), and the surrounding ∼500 bp regions were chosen as homology arms (RHA and LHA). For rhomboid genes, a triple hemagglutinin (3xHA) tag and an additional recodonised sequence of the region spanning the start codon to the FR (RR) was added to the template. The repair templates, LHA:*boxit-*:FR:*boxit+*:RHA (or LHA:3xHA:RR:*boxit-*:FR:*boxit+*:RHA), were fully synthesized commercially (Azenta Life Sciences) for all target genes except for *gap45, gdpd* and *piplc*. These genes were used as test cases to develop modular one-pot assembly protocols for building repair plasmids.

Taking advantage of the common *boxit* sequences, we devised a modular one-pot In-Fusion (TakaraBio) assembly of the repair plasmid. The ∼550 bp long *boxit-*:FR:*boxit+* synthesized segment was amplified using a common primer pair, boxit.F and boxit.R, and the two ∼400 bp long homology arms were amplified from parasite genomic DNA with primers carrying 15 bp extensions. Column-purified amplicons were then assembled into a standard backbone vector in an In-Fusion reaction, according to kit instructions. Alternatively, the same fragments can be assembled by two-step overlap-extension PCR (OE-PCR) using the same primers. Column-purified amplified homology arms and the *boxit-*:FR:*boxit+* containing plasmid (so that the sequence need not be amplified) were added together in the first PCR reaction without any primers. The resulting assembled amplicon was then selectively amplified in the second PCR reaction using outer primers of the homology arms and cloned into the pCR-Blunt vector using Zero Blunt PCR cloning kit (ThermoFisher Scientific). Repair templates produced by OE-PCR were ultimately used for transfections.

The dual-guide targeting plasmid (pCas9-Duo) was designed based on a previously published strategy (*Adikusuma et al., 2017*). The gRNA expression cassette in pDC2-Cas9-gRNA-h*dhfr* (human dihydrofolate reductase)-y*fcu* (yeast cytosine deaminase/uridyl phosphoribosyl transferase) targeting plasmid (*Knuepfer et al., 2017*) was duplicated (gRNA2) but with the ends of the gRNA insertion site modified (TTGG and AAAT in gRNA2 instead of ATTG and CAAA in gRNA1 respectively). To create each targeting plasmid, two gRNA sequences were ordered as oligos with the above-specified overhangs, annealed and inserted into pCas9-Duo in a simple one-pot Golden Gate assembly. The second gRNA cassette is ideally placed near binding site of M13 forward primer, so that successful insertion of gRNA1 and gRNA2 can be confirmed by Sanger sequencing using M13.R and M13.F_R primers respectively.

CloneAmp HiFi PCR Premix (TakaraBio) were used for all PCR reactions. gRNA sequences in the targeting plasmid were confirmed by Sanger sequencing and repair template sequences were confirmed by longread sequencing (Plasmidsaurus). For sequences of oligonucleotides and other synthesized sequences used in this study, please refer to Supplementary Table 1.

### Parasite culture maintenance, synchronisation and transfection

The DiCre-expressing *P. falciparum* B11 line (*Perrin et al., 2018*) was maintained at 37°C in human RBCs in RPMI 1640 containing Albumax II (Thermo Fisher Scientific) supplemented with 2mM L-glutamine. Synchronisation of parasite cultures were done as described previously (*Harris et al., 2005*) by isolating mature schizonts by centrifugation over 70% (v/v) isotonic Percoll (GE Healthcare, Life Sciences) cushions, letting them rupture and invade fresh erythrocytes for 2 h at 100 rpm, followed by removal of residual schizonts by another Percoll separation and sorbitol treatment to finally obtain a highly synchronised preparation of newly invaded ring-stage parasites. To obtain *shiftiko* lines, transfections were performed by introducing 20 μg of targeting plasmid and 60 μg of linearised repair template (purified using Pure-Link™ HiPure Plasmid Midiprep Kit, Invitrogen) into ∽10^8^ Percoll-enriched schizonts by electroporation using an Amaxa 4D Nucleofector X (Lonza), using program FP158 as previously described (*Moon et al., 2013*). Drug selection with 2.5 nM WR99210 was applied 24 hr post-transfection for 4 days with successfully transfected parasites arising at 2-3 weeks. Individual *shiftiko* lines were cryopreserved prior to pooling and starting the SHIFTiKO workflow.

### SHIFTiKO workflow

To create a pooled culture, *shiftiko* lines were treated with sorbitol to roughly synchronise them to ring or young trophozoite stages, their parasitaemia was estimated using a flow cytometer (see Growth and phenotypic assays) and equal number of parasites from each line were added to a fresh culture of RBCs and continuously cultured to attain 5-10% parasitaemia. To induce DiCre-mediated excision in this pool, percoll-synchronized early rings of about 3-5% parasitaemia were treated with rapamycin (10 nM overnight). Mock (DMSO) treated parasites were used as non-induced controls. To obtain mature schizonts at the end of cycle 0, ∼45 h schizonts were percoll-isolated and let to mature by arresting egress using PKG inhibitor 4-[7-[(dimethylamino)methyl]-2-(4-fluorphenyl)imidazo[1,2-α]pyridine-3-yl]pyrimidin-2 amine (compound 2, C2, 1 μM) for 3 h. To allow these parasites to then undergo egress and invasion, egress-stalled schizonts were washed with fresh media to remove C2 and added to fresh erythrocytes (2% haematocrit) and allowed to invade for 4 h with mechanical shaking (100 rpm). Newly formed rings and the remaining non-egressed schizonts were separated using percoll gradient and the rings fraction was further treated with sorbitol to remove residual schizonts. At each time point specified in Figure 3B, culture volumes yielding 20-30 μL packed parasitized RBCs or 10-15 μL isolated schizonts were collected and frozen.

### Barcode sequencing

Genomic DNA was extracted from collected samples using DNeasy Blood and Tissue kit (Qiagen) and DpnI-digested for 1 h. To amplify the barcodes, nested PCR was run with primer pairs barcode.F-barcode.R and padding.F-padding.R using KAPA HiFi HotStart Readymix. Column-purified amplicons were repurified using AMPure beads (Beckman Coulter, 1.8X sample volume). DNA libraries were prepared using Ligation Sequencing Kit (SQK-LSK109) and Native Barcoding Expansion 1-12 (EXP-NBD104) kits, pooled and sequenced either in a MinION R9 flow cell or a Flongle R9 flow cell according to manufacturer’s instructions.

### Growth and phenotypic assays

All assays on individual *shiftiko* lines were performed on uncloned populations. Growth assays were performed to assess parasite growth across 3-4 erythrocytic replication cycles. Synchronous cultures of ring-stage parasites at 0.1% parasitaemia and 2% haematocrit were maintained in triplicates in 48-well plates. 50 μL from each well was sampled at 0, 2, 4, and 6 days post-RAP treatment, fixed with 50 μL of 0.2% glutaraldehyde in PBS and stored at 4°C for flow cytometry quantification. Fixed parasites were stained with SYBR Green (Thermo Fisher Scientific, 1:10,000 dilution) for 20 min at 37°C and analysed by flow cytometry on a BD FACSVerse using BD FACSuite software. For every sample, parasitaemia was estimated by recording 10,000 events and filtering with appropriate forward and side scatter parameters and gating for SYBR Green stain-positive (infected RBCs) and negative RBCs using a 527/32 detector configuration. All data were analysed using FlowJo software.

In the above experiments, samples were also collected 24h post-RAP treatment for running diagnostic PCR on extracted genomic DNA (DNAeasy Blood and Tissue kit, Qiagen) to confirm successful integration and excision. Amplicons from whole locus amplification were gel-extracted and sequenced (Plasmidsaurus).

To assess invasion rates in *rom4-shiftiko* parasites, highly synchronous mature schizonts were added to fresh erythrocytes (2% haematocrit) and allowed to invade for 4 h under either static conditions or with mechanical shaking (100 rpm) (three replicates in each condition). Cultures were sampled before and after the 4 h invasion period and fixed as before for quantification.

### Fluorescence microscopy

For immunofluorescence assays of *rom8-shiftiko* parasites, thin films of parasite cultures containing C2-arrested mature schizonts were air-dried, fixed in 4% (w/v) formaldehyde for 30 min (Agar Scientific Ltd.), permeabilized for 10 minutes in 0.1% (w/v) Triton X-100 and blocked overnight in 3% (w/v) bovine serum albumin in PBS. Slides were probed with anti-MSP1 human monoclonal antibody (X509; 1:1,000 dilution), followed by AlexaFluor 647-conjugated anti-human antibodies (Invitrogen, 1:1,000). Slides were then stained with 1 μg/mL DAPI and mounted in Citifluor (Citifluor Ltd., Canterbury, U.K.).

Z-stacks (125-nm Z-step) were acquired on an VT-iSIM superresolution imaging system (Visitech International), using an Olympus IX83 microscope, 150×/1.45 Apochromat objective (UAPON150XOTIRF), ASI motorized stage with piezo Z, and Prime BSI Express scientific complementary metal oxide semiconductor camera (Teledyne Photometrics). The microscope was controlled with Micro-Manager v2.0 gamma software.

### Data analysis and visualization

Target gene sequences and domain information were downloaded from PlasmoDB database (*Amos et al., 2022*) and floxing strategies and plasmids designed in SnapGene software. To visualise the floxing strategies (Figure 3A), all feature coordinates were plotted using gggenes CRAN package in R.

For barcode-sequencing, basecalling and demultiplexing from raw nanopore output (.fast5 or .pod5 files) was done using Guppy v3.2.2 or Dorado v0.5.0 and the reads mapped onto the 438 bp long amplicon reference sequence using minimap2 v2.2 (-ax map-ont). Exact matches to barcode sequences were counted from the mapped reads.

All statistical analysis and data visualization was performed in R v4.2.2 (*R Core Team, 2021*). Relative growth fitness was calculated by estimating the ratios or change in barcode proportions between Tn and T0 and normalised to changes in barcode proportions of the non-essential control *p230p*. 95% confidence interval of these ratios were derived using delta method of estimating variance. Welch t-test were used to compare group means and where necessary Bonferroni adjustment for multiple comparisons was applied to the p-value of statistical significance.

## ACKNOWLEDGEMENTS

We thank Fiona Hackett and Christine Collins for providing transfection-grade schizonts and Theo Sanderson for valuable tips on bar-seq analysis. This preprint uses a template adapted from Henriques Lab bioRxiv template (C) Ricardo Henriques with some style elements borrowed from eLife template in Overleaf.

## AUTHOR CONTRIBUTIONS

Conceptualization, Funding acquisition: AR and MJB. Data curation, Formal analysis, Investigation, Methodology, Visualization, Writing – original draft : AR. Writing – review & editing: MJB and AR.

## Supplementary Figures

**Figure. S1.**
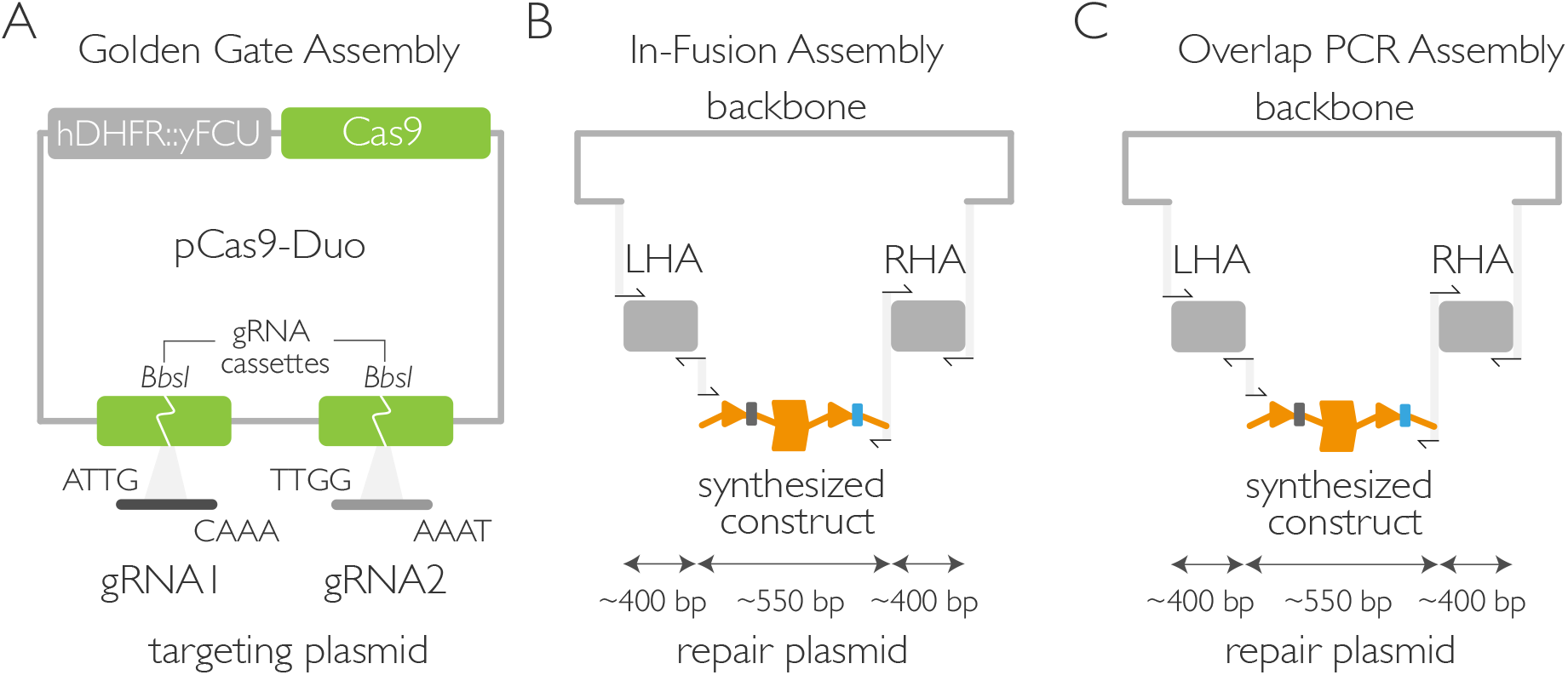
Modular assembly of SHIFTiKO plasmids.

**Figure. S2.**
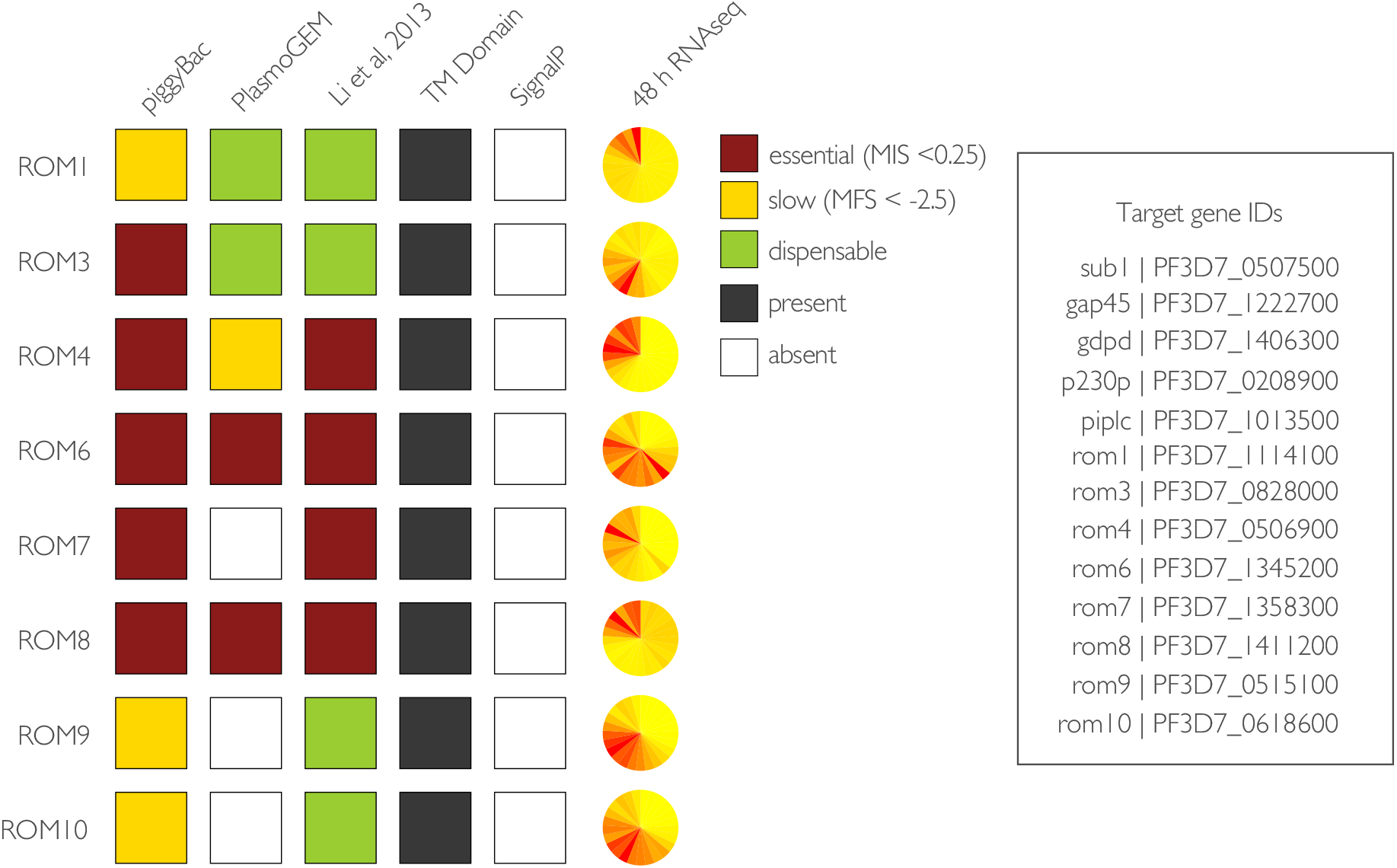
Essentiality of rhomboid proteases in *Plasmodium*.

**Figure. S3.**
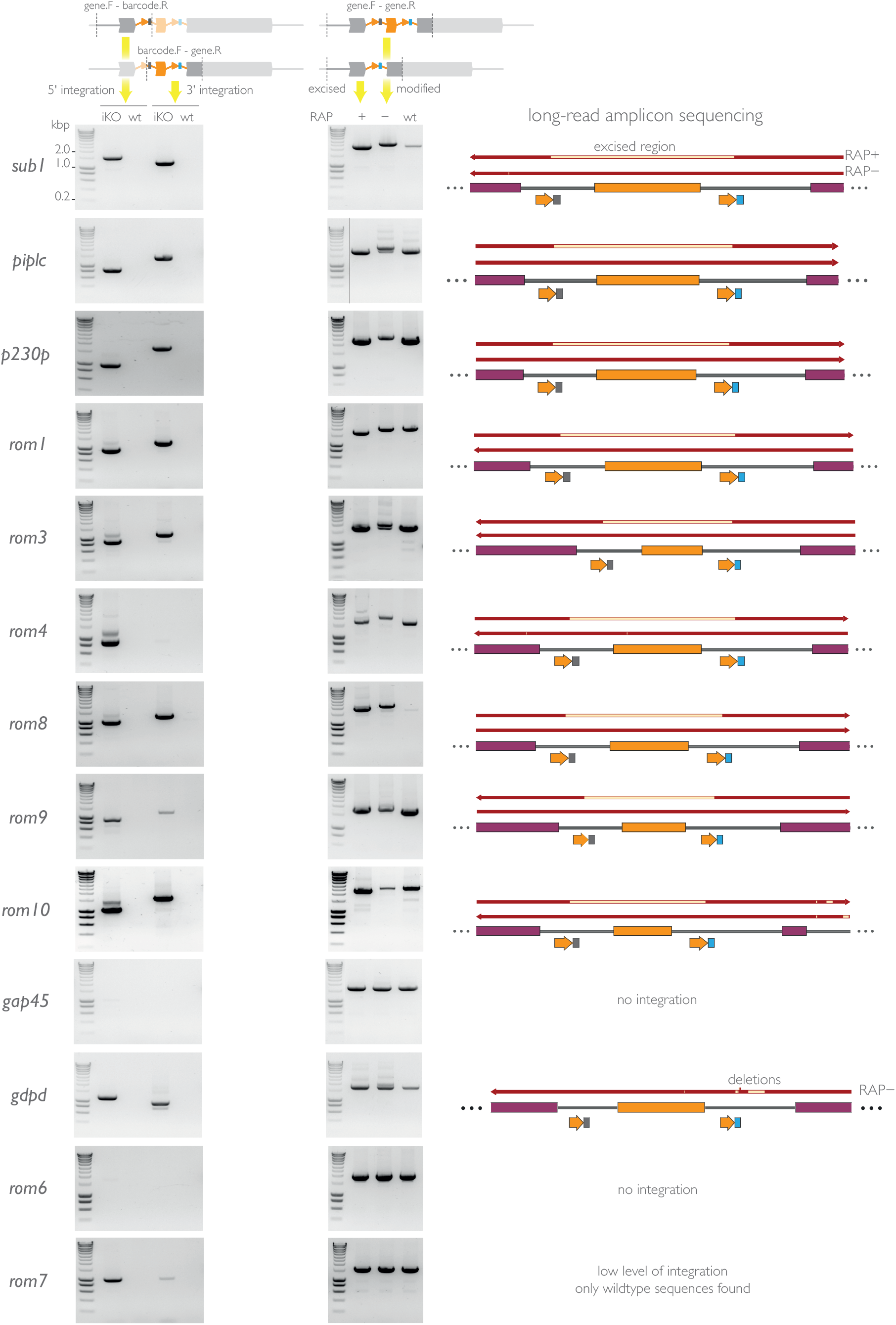
Diagnostic PCR to determine successful integration and excision in individual uncloned *shiftiko* lines.

**Figure. S4.**
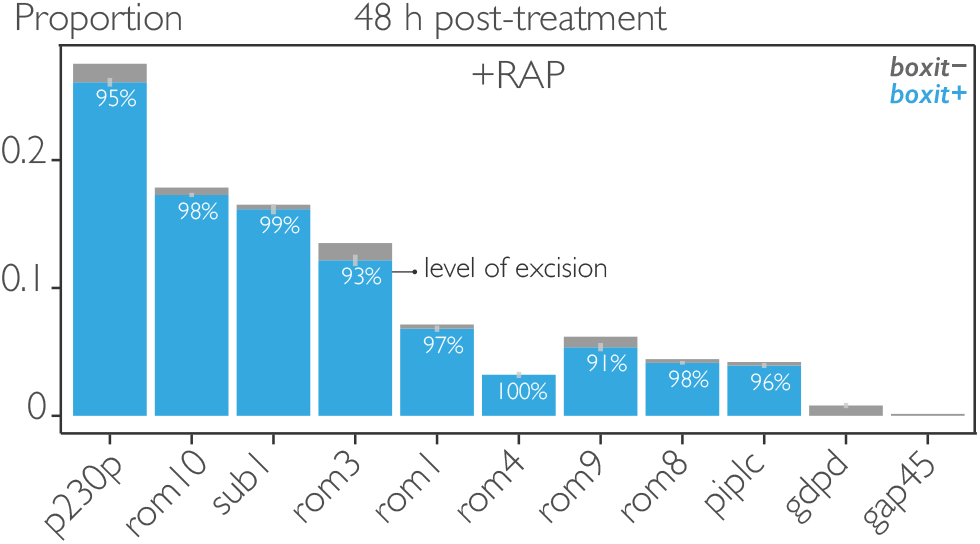
Proportions of *boxit-* and *boxit+* barcodes in mock-(-RAP) and RAP-treated (+RAP) parasites at 48 h post-treatment (T0sz).

**Figure. S5.**
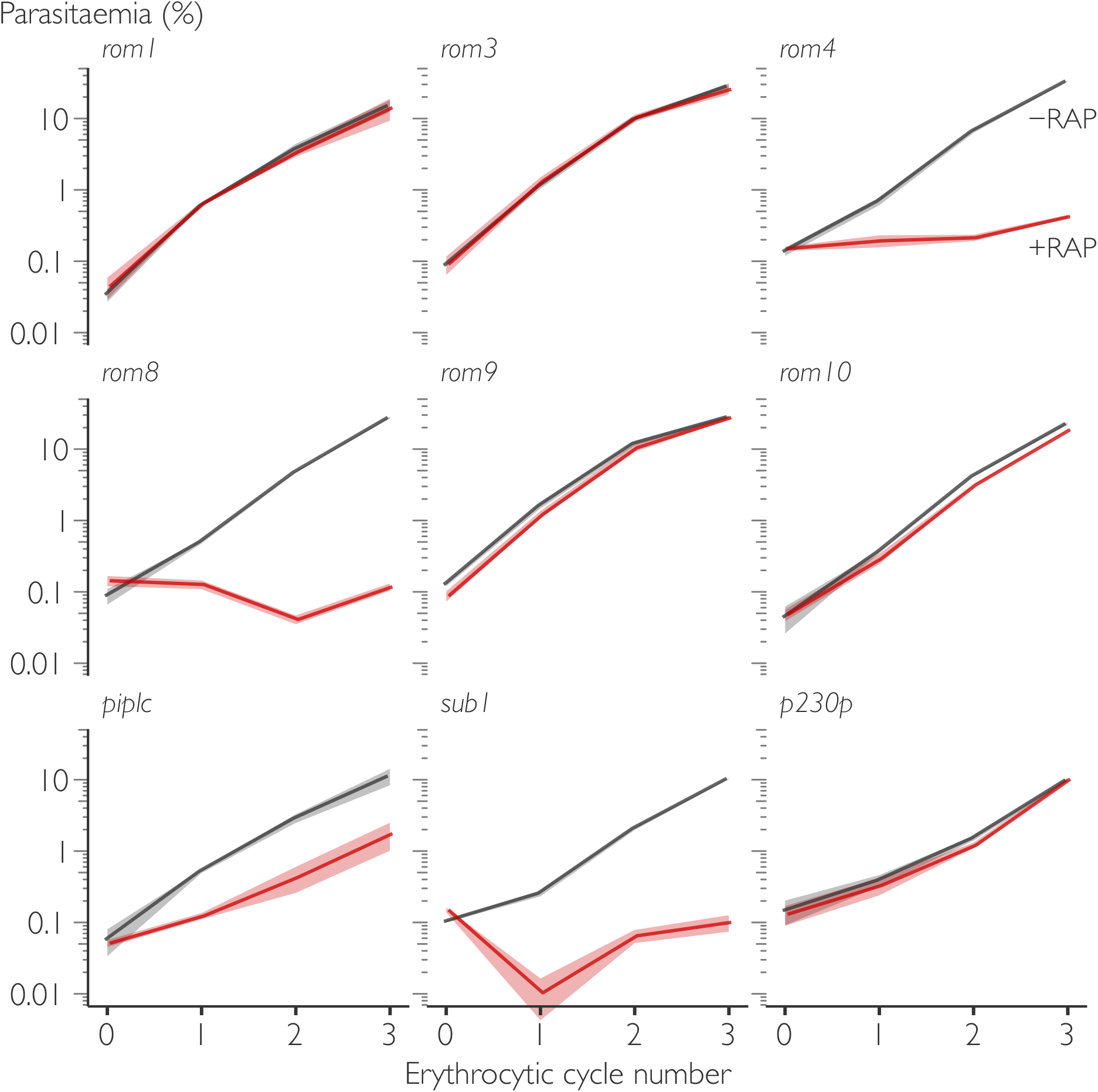
Cell-based growth profiles of mock-(-RAP) and RAP-treated (+RAP) parasites in individual uncloned *shiftiko* lines.

**Figure. S6.**
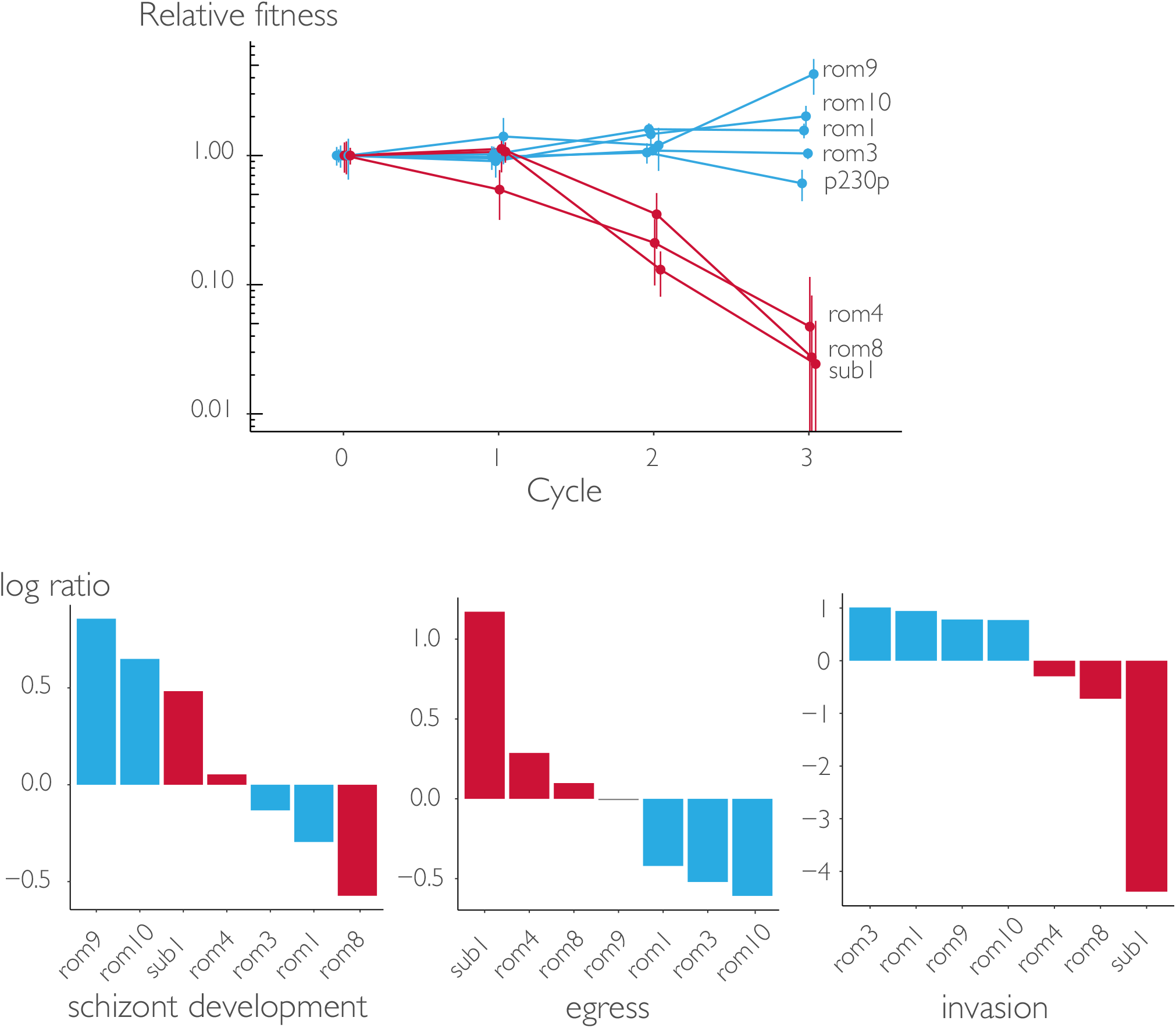
Inducible phenotypic screen results from another independently constituted pool with a subset of the target genes.

